# One predator and two competing prey: insights from a stochastic metapopulation model and mean-field equations

**DOI:** 10.64898/2025.12.12.692652

**Authors:** Daniela Lioren Suárez, María Fabiana Laguna, Nara Guisoni

**Affiliations:** Statistical and Interdisciplinary Physics Group, Centro Atomico, Bariloche (CNEA) and CONICET, R8402AGP Bariloche, Argentina; Universidad Nacional de La Plata (UNLP). La Plata, Buenos Aires, Argentina; Laboratorio de Biología de Sistemas, Centro Regional de Estudios Genómicos, Centro de Endocrinología Experimental y Aplicada, Facultad de Ciencias Exactas-Facultad de Ciencias Médicas, Universidad Nacional de La Plata (UNLP), Consejo Nacional de Investigaciones Científicas y Técnicas (CONICET). La Plata, Buenos Aires, Argentina; Universidad Argentina de la Empresa (UADE). Instituto de Tecnología (INTEC). Buenos Aires, Argentina

## Abstract

We investigate a spatially explicit metapopulation model consisting of one predator and two hierarchically competing prey species on a discrete lattice. Each local population follows stochastic rules for extinction, colonisation, competition, and predation. From the master equation of this individual-based model, we rigorously derive the corresponding mean-field equations. The analysis of these first-principles mean-field equations reveals the existence of a rich phase diagram with different coexistence regimes depending on parameter values. We identify both stable and spiral nodes, in agreement with the damped oscillations observed in Monte Carlo simulations in the latter case. We find qualitative agreement between the mean-field results and Monte Carlo simulations for the three-species coexistence and for the coexistence of the two prey species. However, the mean-field description fails to reproduce the coexistence between the predator and the inferior prey at high predator coverages. We argue that this discrepancy arises from spatial prey aggregation, which the mean-field approach cannot capture since it neglects correlations. In the stochastic model, spatial clustering acts as a crucial protective mechanism against predation, particularly for the best coloniser. Our findings suggest that prey aggregation contributes to system stability when colonisation and predation operate at comparable spatial scales. The combination of first-principles mean-field equations and stochastic simulations constitutes a powerful framework for clarifying the roles of hierarchical interactions, predation, colonisation, spatial organisation and stochasticity in multi-species coexistence.

## I. INTRODUCTION

The use of quantitative models in ecology is fully extended [1]. Historically, the most common approach has been to describe the system at the population level through differential equations. These models can be constructed without providing a detailed description of the individuals or species involved and how they interact. Instead, they focus on capturing observed patterns and empirical relationships between them, where the terms in the governing equations represent the overall effects of the interactions present in the system. In such phenomenological models, the functional forms of the equations are proposed *ad hoc*, without asking about the underlying fundamental mechanisms that generate the observed phenomena.

Paradigmatic examples of the *ad hoc* approach include the Levins’ metapopulation model [2] and the Lotka-Volterra equations [3]. More complex predator–prey models based on differential equations have also been extensively developed. These can incorporate hierarchical competition between species to study, for instance, habitat destruction [4, 5] or global extinction through synchronization [6]. Recent reviews have further explored the diversity of applications of predator–prey models, highlighting their development in theoretical and applied studies [7, 8]. Phenomenological models have also been applied to biological control, exploring scenarios such as alternative food sources, intra-guild predation, and lifestage structure [9–11].

An alternative approach in ecological modelling relies on individual-based models. Here, a finite number of individuals, each characterized by a small set of attributes, interact according to predefined probabilistic rules. Prior biological knowledge about the individuals or species and their interactions should be considered. The temporal evolution of these models follow a Markov process, and space is usually represented explicitly. Individual-based models can be studied exactly through Monte Carlo simulations, which allow the exploration of stochastic effects and spatial correlations. Numerous examples exist, including the Levin and Segel model [12], the stochastic metapopulation models developed in [13], and different versions of cyclic predator–prey models [14–18]. Comparisons between individual-based predator–prey models and their *ad hoc* mean-field counterparts have been carried out for systems of two [19, 20] and three species [4]. Qualitative agreement between the results is not always achieved [4, 20].

Individual-based models can also be approached analytically through the master equation formalism [21, 22]. The master equation governs the time evolution of the probability distribution of all possible system configurations and provides an exact stochastic description of the system [23]. From the master equation, it is possible to derive the corresponding differential equation-based model analytically from an individual-based model, via mean field or by considering different approximations for correlations [1]. Unlike the *ad hoc* models mentioned previously, the differential equations in this case are derived from first principles and a well-defined approximation scheme is employed. Consequently, the resulting meanfield equations are unambiguous and, within the limits of the approximation used, provide an accurate macroscopic description of the system.

Several authors have pursued this approach. In a seminal work, McKane and Newman [21] compared meanfield equations derived from first principles with standard *ad hoc* mean-field equations commonly found in population biology textbooks. They demonstrated that even for very simple models, important differences may arise, including the appearance of cross-diffusion terms that cannot be introduced *ad hoc*. Satulovsky and Tomé [24] proposed a stochastic metapopulation (or lattice-gas) model for a two-species predator–prey system. The model included local interactions analogous to those of the contact process. By applying different approximation schemes for spatial correlations, they derived first-principles meanfield equations and identified a rich phase diagram. This diagram featured multiple absorbing states and a region with oscillatory predator–prey dynamics, in qualitative agreement with Monte Carlo simulations. Later, Satulovsky [25] derived mean-field equations containing negative cross-diffusion terms from the master equations of different versions of lattice-based Lotka–Volterra models. Satulovsky found that the mean-field equations could not reproduce the spatial patterns that emerged in the stochastic models for certain parameter values. In fact, qualitative agreement between stochastic and first-principle mean-field models is not always achieved, particularly when spatial correlations and stochastic effects are important. In this cases, more refined mean-field approximations are needed [1, 17, 18, 21, 26].

In this work, we study a spatially explicit three-species metapopulation model consisting of one predator and two prey species. The model incorporates the biological processes of extinction, interspecific competition between preys, predation, and colonisation. Special colonisation rules account for the predator’s search for prey and for hierarchical interspecific competition between prey. A similar version of this model was previously studied by one of us and collaborators through stochastic simulations and *ad hoc* mean-field equations [4]. Here, we propose a first-principles derivation of the mean-field equations. This involves writing the master equation for the model and obtaining a mean-field approximation of the time evolution of the mean value of each species. We contrast our formulation with that earlier phenomenological approach. Our first-principles framework provides a welldefined link between the individual-based stochastic rules and the resulting population-level dynamics, allowing us to determine analytically how the parameters used in both descriptions relate to each other. Our derivation not only clarifies the origin and interpretation of the terms appearing in the mean-field equations, but also facilitates a systematic comparison with Monte Carlo simulations. In this sense, the present work helps bridge the gap between phenomenological and first-principles population-level formulations in ecological modelling.

We also perform a steady-state analysis of the first-principles mean-field equations and study the stability of the stationary solutions, showing that the model exhibits a rich phase diagram with different coexistence regimes depending on parameter values. We also carry out stochastic simulations, finding qualitative agreement with our first-principles mean-field results regarding the coexistence of three species and of the two prey species. However, the mean-field equations do not accurately describe the coexistence of the predator and inferior prey species that occurs for high predator occupancy. We argue that, in this situation, the spatial aggregation of inferior prey plays a crucial role that is not captured by mean-field equations. The combination of first-principles mean-field equations and stochastic simulations offers a deeper understanding of how hierarchical interactions, predation, colonisation, spatial prey aggregation, and stochasticity, shape the coexistence of species.

The paper is organized as follows. In Section II we motivate and define the model rules. In Section III we derive the master equation. In Section IV we propose a mean-field approximation for the master equation, make a comparison with a previous *ad-hoc* mean-field model, present a stability analysis of the stationary states and show results for the numerical solutions of the mean-field equations. In Section V we present stochastic simulations of the model, and compare these results with mean-field equations. Conclusions are given in Section VI.

## II. THE THREE-SPECIES METAPOPULATION MODEL

We consider a spatially explicit three-species metapopulation model originally proposed by Laguna *et al*. [4]. The system consists of two prey species, denoted by *X* and *Y*, which engage in hierarchical competition, with *X* being the superior (dominant) species and *Y* the inferior one. A third species, *Z*, preys on both.

Each species forms an assemblage that is spatially distributed over a two-dimensional regular lattice composed of Ω = *L* × *L* habitat patches of equal area. In metapopulation models, the focus is on determining whether a species is present or absent in a patch, rather than how many individuals inhabit it. Patches can be simultaneously occupied by one, two, or all three species.

In the following, *X*_*i*_, *Y*_*i*_, and *Z*_*i*_ indicate that patch *i* is occupied by species *X, Y*, and *Z*, respectively. Conversely, 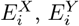, and 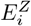 denote that patch *i* is unoccupied by the corresponding species. The species interact through four biological processes: extinction, interspecific competition between prey, predation, and colonisation. Interspecific competition between the two prey species may arise, for example, from differences in body size or resource acquisition efficiency. These processes determine the spatio-temporal dynamics of occupied and unoccupied patches, according to the following rules:

i. *Extinction*: any species occupying a patch *i* can go extinct with probability *e* (*e*_*x*_ for *X, e*_*y*_ for *Y* and *e*_*z*_ for *Z*), thereby vacating the patch.

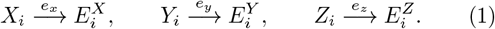
ii. *Interspecific competition between preys*: when species *X* and *Y* inhabit the same patch *i, X* can competitively exclude *Y* from the patch with probability *c*. Here, *X* is the superior competitor.

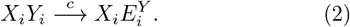
iii. *Predation*: if the predator *Z* shares a patch *i* with either prey, it can prey upon them with probability *µ* (*µ*_*x*_ for *X* and *µ*_*y*_ for *Y*).

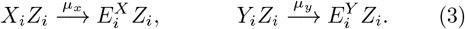
iv. *Colonisation*: each of the three species can colonize neighbouring sites with a certain probability (*c*_*x*_ for *X, c*_*y*_ for *Y* and *c*_*z*_ for *Z*), provided specific conditions are satisfied.

- Species *X* occupying a patch *i* can colonize a neighbouring patch *j* that is unoccupied by its own species:

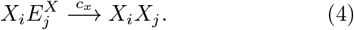
- Species *Y* occupying a patch *i* can colonize a neighbouring patch *j* that is unoccupied by itself and by the superior competitor *X*:

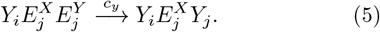
- Species *Z* occupying a patch *i* can colonize a neighbouring patch *j* that is unoccupied by its own species but occupied by at least one of the prey species. Therefore, three possible configurations of prey occupation in the target patch allow for *Z* colonisation, depending on whether the patch contains *X, Y* or both:

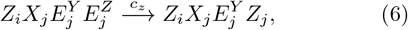

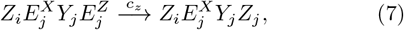

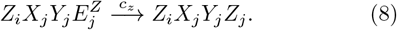

Note that the colonisation process, although it involves two neighbouring patches, differs from migration, since the latter entails the movement of a species from one patch to another (vacating one site while occupying the next) whereas colonisation results in the occupation of an additional site. For the purposes of the dynamical model, colonisation is therefore equivalent to birth.

The time evolution of the model can now be described. The biological processes considered are in three distinct situations: extinction, which involves sampling one patch and one species; hierarchical competition and predation, which require sampling one patch and two species; and colonisation, which involves two neighbouring patches and one species. At each time step we sample a patch *i* with probability 1*/*Ω. After that, to ensure that all processes occur with the appropriate probability, the following situations are considered:

i. On a fraction *q*_1_ of the occasions we randomly choose a species from patch *i* with probability 1*/*3. If the patch is not inhabited by the chosen species, nothing happens. Otherwise, extinction may occur according to Eqs.1.
ii. On a fraction *q*_2_ of the occasions we randomly choose two species from patch *i* with probability 1*/*6. If the patch is unoccupied by one or both species, nothing happens. Otherwise, if the two species are preys (that is *X*_*i*_ and *Y*_*i*_), hierarchical competition can occur according to Eq. 2. And, in case we choose the predator (*Z*_*i*_) and one prey (*X*_*i*_ or *Y*_*i*_), predation may occur according to Eqs. 3.
iii. Finally, for a fraction 1 − *q*_1_ − *q*_2_, another patch *j* is chosen at first neighbours of patch *i* with probability 1*/q*, where *q* refers to the coordination number, equal to 4 here. One species is drawn from patch *i* with probability 1*/*3. If the patch is not inhabited by the chosen species, nothing happens. Otherwise, colonisation can occur according to Eqs. 4-8, depending on the species sorted.

The present model is a slight modification of a model previously proposed by one of us and collaborators [4]. The difference lies in how time evolution is implemented. In [4], after sampling a patch, each species was sequentially tested for the probabilistic occurrence of colonisation, extinction, predation, and interspecific competition. In contrast, the current implementation introduces two random parameters, *q*_1_ and *q*_2_, ensuring that all biological processes are treated as fully stochastic events: extinction occurs with probability *q*_1_, competition and predation with probability *q*_2_, and colonisation with probability 1 − *q*_1_ − *q*_2_. In the present implementation, no event occurs deterministically or in sequence; all are probabilistic.

## III. THE MASTER EQUATION

The stochastic rules described above can be formally expressed in terms of a master equation, which governs the time evolution of the probability distribution of all possible system configurations. This formulation provides the analytical framework linking the individual-level processes to the macroscopic, population-level dynamics, and establishes the connection between stochastic simulations and their first-principle mean-field representations.

We will define 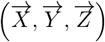 as the state of the system, given that 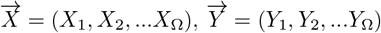 and 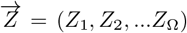, remembering that Ω = *L* × *L* is the number of patches. Let the occupation state of species *X* in patch *i* be given by the binary variable *X*_*i*_ ∈ {0, 1}, where *X*_*i*_ = 1 denotes occupation and *X*_*i*_ = 0 denotes non-occupation. Similarly, *Y*_*i*_ ∈ {0, 1} and *Z*_*i*_ ∈ {0, 1} indicate the occupation states for species *Y* and *Z*, respectively. Let 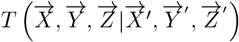 be the transition probability from state 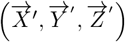 to state 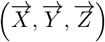. Since only metapopulation transitions from 0 → 1 and from 1 → 0 may take place, the only nonzero transition probabilities of the processes given by Eqs. 1-8 are:

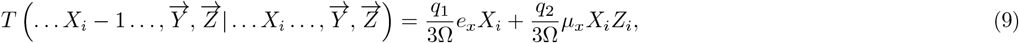

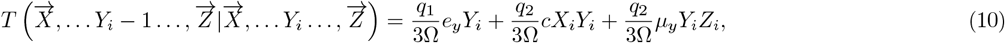

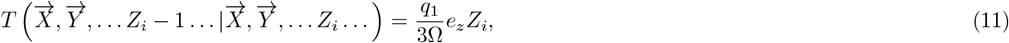

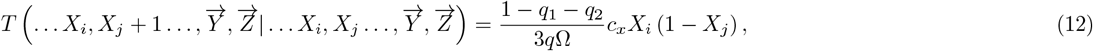

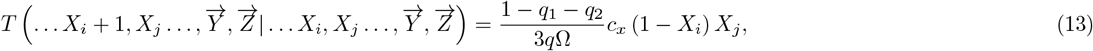

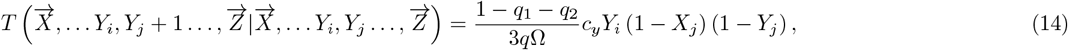

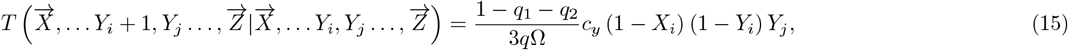

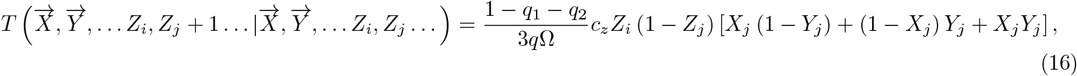

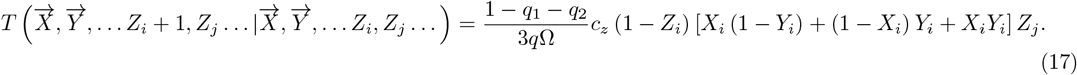

For simplicity, only the occupations of the patches where changes occur (*i* or *j*) have been explicitly shown on the left-hand side of the Eqs. 9-17. Metapopulation vanishing (*X, Y, Z* = 1 → 0) is governed by extinction, interspecific competition, and predation (Eqs. 9-11); while metapopulation appearance (*X, Y, Z* = 0 → 1) is conducted by colonisation (Eqs. 12-17). As species can colonise from patch *i* to *j*, or vice versa, there are two equations for the transition probabilities of each species. One is related to the appearance of a metapopulation in patch *i*, and the other is connected to the appearance of a metapopulation in patch *j*.

Eqs. 9-17 are one-step Markov process, so we can immediately write down a master equation describing how the probability of finding the system in the state 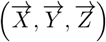 at a given time *τ*, that is 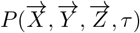, changes over time:

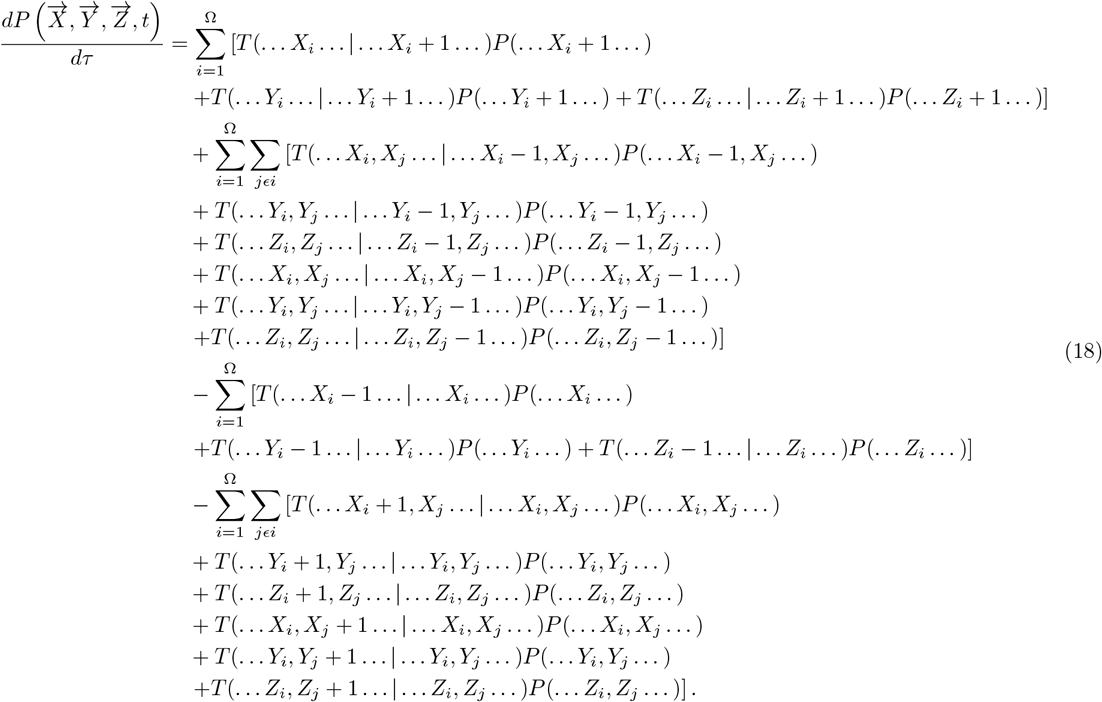

Note that the master equation is simply the sum of transitions from states with *X*_*i*_ ±1, *Y*_*i*_ ±1, or *Z*_*i*_ ±1 to the target state 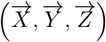, minus the sum of transitions from the target state 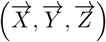 to states with *X*_*i*_ ± 1, *Y*_*i*_ ± 1, or *Z*_*i*_ ± 1. To consider all possible configurations, processes involving one patch, that is extinction, hierarchical competition and predation, must be summed over all Ω patches. For colonisation, an additional sum must be made over the *j* neighbours of each site *i*, despite the sum already being made over all Ω patches. The notation *jϵi* represents the sum of all sites *j* that are nearest neighbours of site *i*.

The master equation of the system (Eq. 18) allows us to calculate the time evolution of the mean value of each species in a given patch *k*. For example, for species *X*, we have:

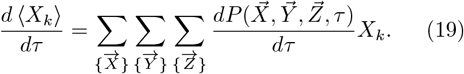

Analogous expressions are obtained for *Y* and *Z*.

Substituting Eqs. 18 and 9-17 in Eq. 19, making variable changes of the form 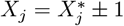 (and similarly for the other variables), and finally using the symmetries of the intermediate equations, we found the following equations for each species. Similar procedure is employed for *Y* and *Z*.

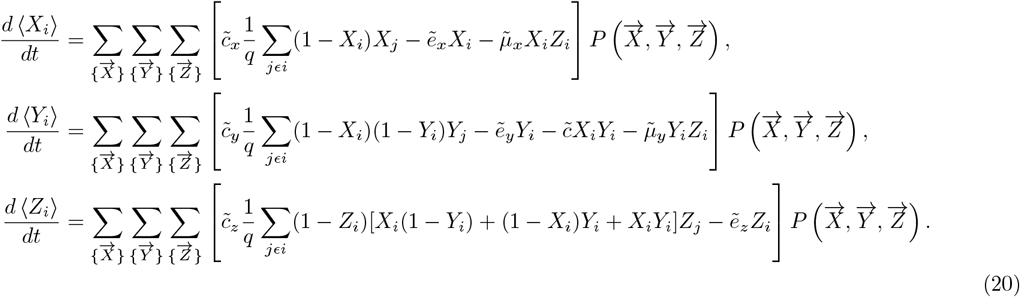

where, for simplicity, we define new quantities as follows:

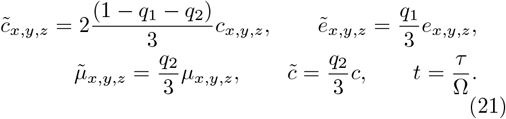

Now, simply using the definition of the mean value in Eqs. 20 we can derive the following equations:

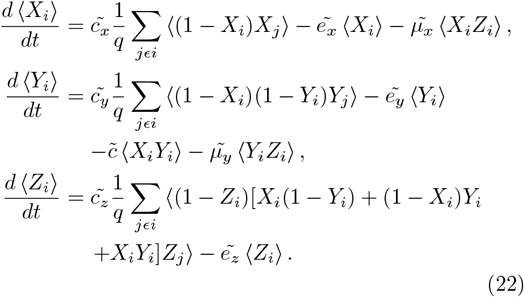

## IV. MEAN-FIELD APPROXIMATION

To obtain approximate analytical solutions of Eqs. 22, we apply a truncation scheme. In its simplest version, this consists of expressing the mean value of a cluster of patches or species as the product of the mean values of each component. For example, ⟨*X*_*i*_*X*_*j*_⟩= ⟨*X*_*i*_⟩ ⟨*X*_*j*_⟩ or ⟨*X*_*i*_*Y*_*i*_*Z*_*i*_⟩ = ⟨*X*_*i*_⟩ ⟨*Y*_*i*_⟩ ⟨*Z*_*i*_⟩. This approximation neglects correlations between patch occupancy and species co-occurrence. Under this assumption, Eqs. 22 become

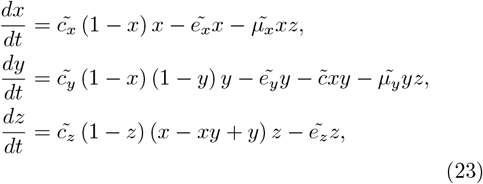

where ⟨*X*_*i*_⟩ = ⟨*X*_*j*_⟩ = ⟨*X*⟩ = *x*, ⟨*Y*_*i*_⟩ = ⟨*Y*_*j*_⟩ = ⟨*Y* ⟩ = *y*, and ⟨*Z*_*i*_⟩ = ⟨*Z*_*j*_⟩ = ⟨*Z*⟩ = *z*, since we are assuming spatial homogeneity and thus seeking homogeneous solutions. The variables *x, y*, and *z* can be interpreted as the densities of species *X, Y*, and *Z*, respectively. Additionally, we have considered the thermodynamic limit Ω → ∞. The relationship between the parameters of the stochastic model and the mean-field parameters is univocally defined by Eq. 21.

Equations 23 describe the temporal dynamics of the average metapopulation occupancy of species *X, Y*, and *Z* within the mean-field approximation. In fact, such equations could be naively written down on an *ad hoc* basis. The first term in each equation corresponds to colonisation and is proportional to 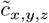. These terms depend on the species densities and the densities of available patches satisfying the conditions specified in Eqs. 4–8. In Eq. (23) for *Y*, the term 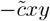 represents interspecific competition, leading to a decrease in the density of the inferior species. The terms 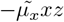 and 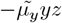 describe predation by the predator *Z* on each prey. Each species also has an extinction term proportional to its respective probability *e*_*x,y,z*_.

Following the definition of the Laplacian operator, different mean-field equations can be derived from Eqs. 22, as made previously [24]. Notably, this approximation gives rise to cross- and self-diffusion terms in the meanfield equations as shown in Appendix A. For the parameter values considered here, the steady-state results obtained from this modified mean-field version coincide with those of Eqs. 23. Therefore, we restrict our analysis to the simpler mean-field formulation. Similar results were reached by Satulovsky for three versions of simple predator–prey models [24], showing that spatial instabilities do not emerge from these first-principles mean-field equations that include cross- and self-diffusion terms.

### IV.1. Comparison with an *ad hoc* mean-field model

A phenomenological mean-field version of the present model were previously proposed by one of us and collaborators [4]. However, this formulation differs from the present one in the colonisation terms of *Y* and *Z*, as we discuss next.

The colonisation term of the superior prey is identical in both approaches, since in each case the growth of species *X* depends only on the availability of empty sites and on its own occupancy through a factor proportional to (1 − *x*)*x*. The first relevant difference appears in the equation for the inferior prey. In the *ad hoc* model, colonisation of species *Y* depends on the fraction of sites not occupied by either prey and on its own occupancy through the factor (1 − *x* − *y*)*y*, whereas in the first-principles formulation it takes the form (1 − *x*)(1 − *y*)*y*. This expression incorporates explicitly the outcome of hierarchical competition and introduces an additional nonlinear contribution. Although competition between the two prey is represented in both models by a loss term proportional to *xy*, the altered structure of the colonisation process modifies the effective growth of *y*, thereby changing the net impact of competitive interactions.

A second difference concerns the predator colonisation term. In the *ad hoc* formulation, predator growth is proportional to (*x* − *xy* + *y*)*z*, whereas in the first-principles version it takes the form (1 − *z*)(*x* − *xy* + *y*)*z*, reflecting the explicit requirement that the predator can colonise only those patches that are not already occupied by its own species. This additional factor not only modifies the form of predator colonisation, but also introduces further non-linearities that affect predator persistence and the boundaries of coexistence.

These structural differences illustrate how the first-principles approach provides a more systematic and transparent connection between the individual based stochastic dynamics and their population level representation. As a consequence, the first-principles formulation captures non-linearities that are often difficult to incorporate correctly in phenomenological models, particularly in systems such as the one analysed here, where complex colonisation rules can make *ad hoc* equations challenging to construct.

Another important difference between our formulation and phenomenological proposals concerns the relationship between stochastic model parameters and mean-field parameters. These can only be determined unambiguously within first-principles approaches, as presented here (see Eq. 21).

### IV.2. Stationary states and stability analysis

To study the steady-state solutions of Eqs. 23 and their stability, it is convenient to rewrite them as

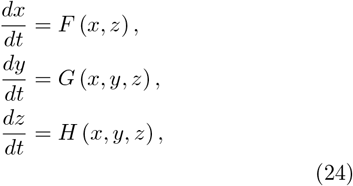

where 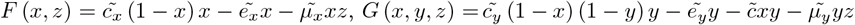 and 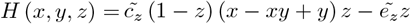.

The steady-state solutions of Eqs. 24, *S*_*i*_ = (*x*_*ss*_, *y*_*ss*_, *z*_*ss*_), are the positive solutions of *F* (*x, z*) = 0, *G* (*x, y, z*) = 0, and *H* (*x, y, z*) = 0. Depending on the parameter values the system can present up to eight fixed points: the trivial solution *S*_0_ = (0, 0, 0); the three-species coexistence solution *S*_1_ = (*x*_*ss*_ *>* 0, *y*_*ss*_ *>* 0, *z*_*ss*_ *>* 0); three solutions with coexistence of two species: *S*_2_ = (0, *y*_*ss*_ *>* 0, *z*_*ss*_ *>* 0), *S*_3_ = (*x*_*ss*_ *>* 0, 0, *z*_*ss*_ *>* 0), and *S*_4_ = (*x*_*ss*_ *>* 0, *y*_*ss*_ *>* 0, 0); and three single-species solutions *S*_5_ = (*x*_*ss*_ *>* 0, 0, 0), *S*_6_ = (0, *y*_*ss*_ *>* 0, 0), and *S*_7_ = (0, 0, *z*_*ss*_ *>* 0). In order to simplify notation, from now on, we identify the steady states in following way: *S*_1_ = *XY Z, S*_2_ = *Y Z, S*_3_ = *XZ, S*_4_ = *XY*, *S*_5_ = *X, S*_6_ = *Y*, and *S*_7_ = *Z*.

Linearising the system around each steady state *S*_*i*_ = (*x*_*ss*_, *y*_*ss*_, *z*_*ss*_) allows us to determine its stability. The Jacobian matrix *J* associated with Eqs. (24) is given by

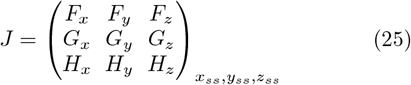

where 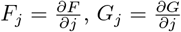 and 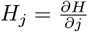, with *j* = *x, y, z* as defined in Appendix B (Eq. 30).

The corresponding secular equation is straightforward

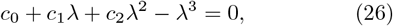

where the exact coefficients *c*_0_, *c*_1_, and *c*_2_ are given explicitly in Appendix B (Eq. 31). The solutions of this cubic equation yield the eigenvalues *λ*, whose analysis determine the stability of each steady state.

The eigenvalue analysis was performed both analytically and numerically for the eight steady states *S*_*i*_, considering the parameters values of interest in the present work (more details in the next section). We found that the system is always monostable. However, different stationary-state solutions can alternate as a control parameter varies. In particular, increasing *e*_*z*_ induces a sequence of transitions: from a two-species coexistence state (*XZ* steady-state), to a three-species coexistence state (*XY Z* steady-state), and finally to another two-species coexistence state (*XY* steady-state), as illustrated in Fig. 2. We considered *e*_*z*_ *>* 0, since there are infinite steady-state solutions of Eqs. 24 for *e*_*z*_ = 0.

**FIG. 1.**
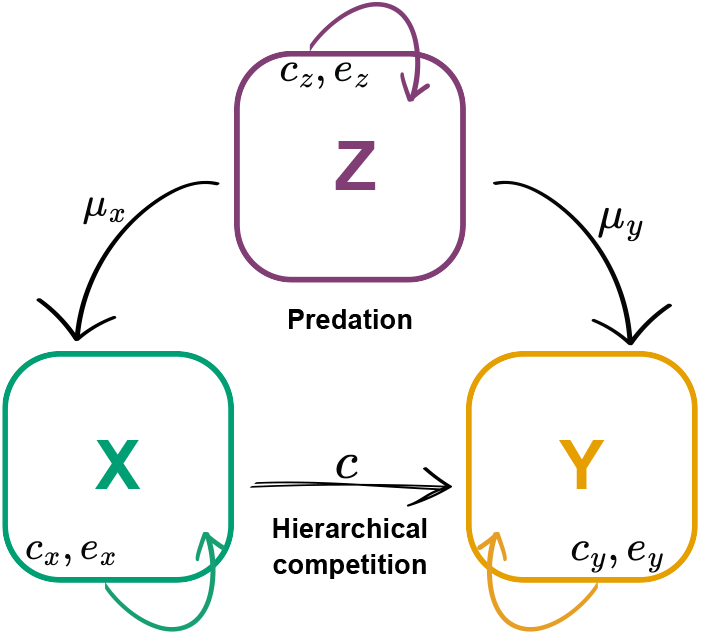
Conceptual model of species interactions: X and Y are prey and Z is a predator. Arrows depict interaction directions; the arrow from X to Y highlights the asymmetry in hierarchical competition.

**FIG. 2.**
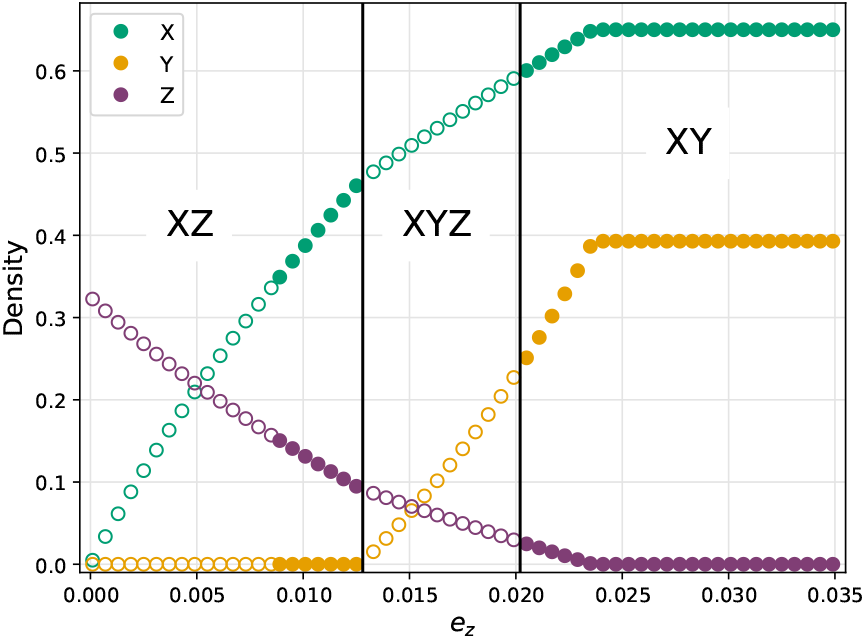
Stationary densities of the species *X* (superior prey, green), *Y* (inferior prey, yellow) and *Z* (predator, purple), as a function of predator extinction probability *e*_*z*_, classified by fixed-point stability. Empty symbols denote spiral nodes and filled symbols stable nodes. Solutions alternate between two- and three-species coexistence states. Parameters values according to Eqs. 21 and *q*_1_ = *q*_2_ = 1*/*3, *c*_*x*_ = 0.05, *c*_*y*_ = 0.1, *c*_*z*_ = 0.015 *e*_*x*_ = 0.035, *e*_*y*_ = 0.01, *µ*_*x*_ = 0.2, *µ*_*y*_ = 0.8, *c* = 0.05.

The stable fixed points can be classified as either stable nodes, when all eigenvalues *λ* are real and negative, or spiral nodes, when two eigenvalues are complex conjugates with negative real parts and the third is real and negative. Both types of stability are shown in Fig. 2.

Numerical solutions of polynomial equations (steady-state solutions of Eqs. 24 and solution of Eq. 26) were obtained numerically using the Newton–Raphson method, as implemented via a custom Mathematica script.

### IV.3. Mean-field numerical solutions

To numerically solve the mean-field equations (Eqs. 23), we used a second-order Runge–Kutta method with a time step of 0.01, which provided good convergence. To ensure the reproducibility of results, we set an arbitrary initial condition *x*(*t* = 0) = *y*(*t* = 0) = *z*(*t* = 0) = 0.5, unless stated otherwise.

We used the parameters values defined in Table I. These follow those proposed in [4], except for the value of *e*_*x*_. For simplicity, we set *q*_1_ = *q*_2_ = 1*/*3. Intraspecific competition favours the superior prey (*X*) over the inferior prey (*Y*). According to the parameters considered here the inferior prey is more likely to be preyed upon (*µ*_*y*_ *> µ*_*x*_), while the superior prey has a higher local extinction probability (*e*_*x*_ *> e*_*y*_). The inferior prey is a better coloniser than the superior prey (*c*_*y*_ *> c*_*x*_), except in a region of Fig. 4. The predator is the weakest coloniser (*c*_*z*_ *< c*_*x,y*_). Following [4], we analyse the effect of varying the predator extinction probability *e*_*z*_. In addition, we explore the system’s response to vary *c*_*y*_.

**TABLE 1.** Parameter values for the three-species metapopulation model. For parameters with a range, the values in brackets indicate the interval explored.

| Parameter | Symbol | Value |
| --- | --- | --- |
| $X$ colonisation | $c_x$ | 0.05 |
| $Y$ colonisation | $c_y$ | 0.1, [0, 0.5] |
| $Z$ colonisation | $c_z$ | 0.015 |
| $X$ extinction | $e_x$ | 0.035 |
| $Y$ extinction | $e_y$ | 0.01 |
| $Z$ extinction | $e_z$ | [0, 0.03] |
| $X$ predation | $\mu_x$ | 0.2 |
| $Y$ predation | $\mu_y$ | 0.8 |
| $XY$ competition | $c$ | 0.05 |

Figure 3 shows the time evolution of the densities of species *X, Y*, and *Z* for two different values of the predator extinction probability, *e*_*z*_, and two different initial conditions. In both panels, in the steady state, only the predator and the superior prey coexist. Increasing *e*_*z*_ results in a higher density of patches occupied by prey relative to the predator. The system exhibits a transient period before reaching the steady state, as evident from the different initial conditions considered. Damped oscillations, which are a typical signature of spiral-node stability, are clearly observed in the upper panel. Oscillations are not expected in the lower panel since, for this value of *e*_*z*_, the fixed point corresponds to a stable node. These different types of stable fixed points were identified through the stability analysis presented in the previous section and illustrated in Fig. 2. In the region of spiral-node fixed points where *e*_*z*_ *<* 0.009, we observed that oscillations became more pronounced as the probability of predator extinction decreased. Similar behaviour in relation to the probability of predator extinction was reported in a simpler predator-prey model exhibiting true oscillations [24].

**FIG. 3.**
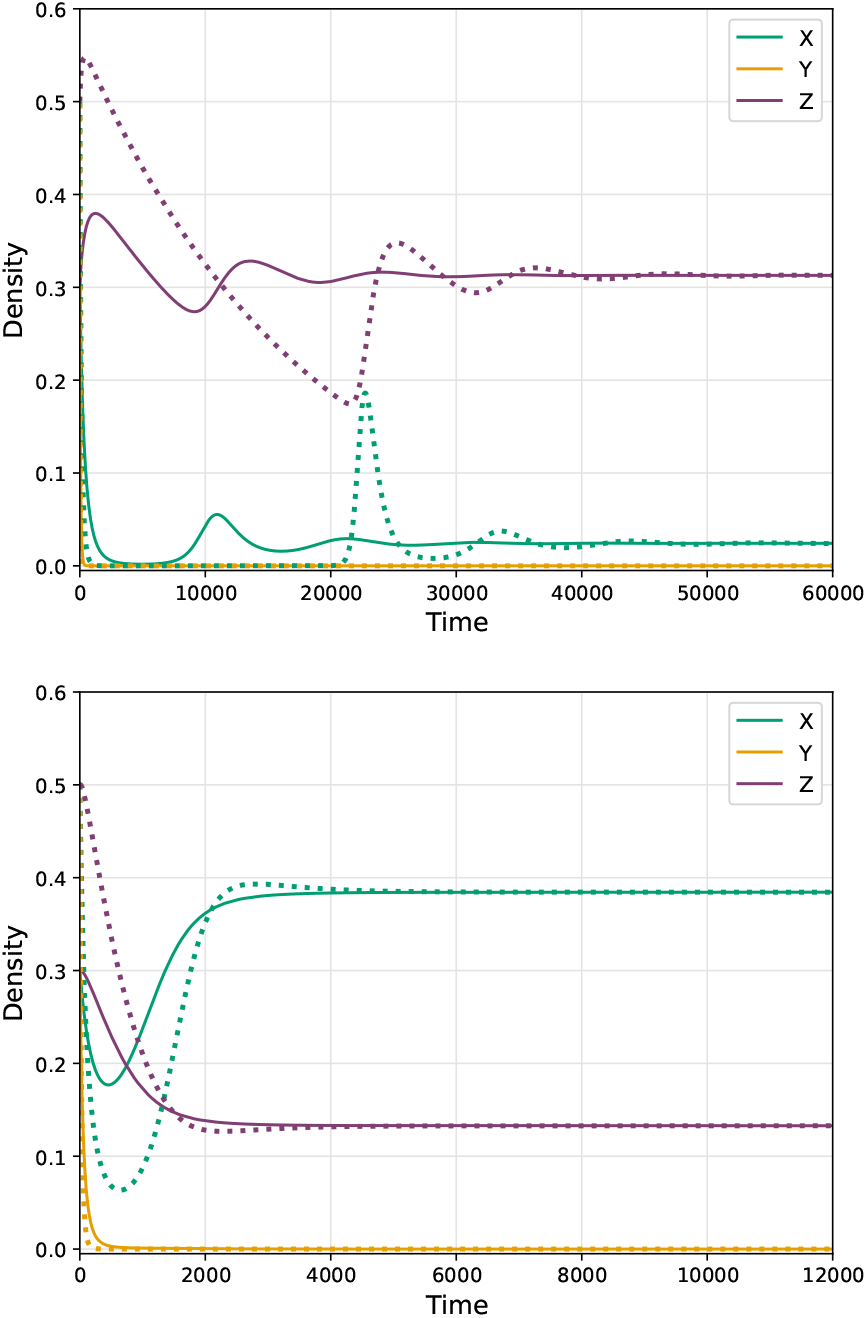
Mean-field results for the time evolution of the densities of species *X, Y* and *Z*. The parameters are listed in Table I, with *c*_*y*_ = 0.10 for both cases, and *e*_*z*_ = 0.0005 in the upper panel and *e*_*z*_ = 0.01 in the lower one. Solid lines correspond to initial densities of 0.3 for all species, and dotted lines to initial densities of 0.5 for all species.

Let us make a more in-depth analysis of Fig. 2 and examine the different steady states exhibited by the system as the probability of predator extinction varies. For low values of *e*_*z*_, the predator and the superior prey coexist, whereas the inferior prey goes extinct (*XZ* steady state). For intermediate *e*_*z*_ values (0.013 *< e*_*z*_ *<* 0.024), all three species coexist (*XY Z* steady state), and when the predator becomes extinct, the system reaches a two-prey coexistence state (*XY* steady state). Across the entire *e*_*z*_ range, the superior prey maintains a higher density of occupied patches than the inferior prey. These results suggest that the predator limits prey abundance, particularly that of the inferior prey, and that the superior prey exerts strong competitive control over the inferior one.

Following the analysis of the steady-states, we now explore how the balance between species depends on the prey colonisation abilities and predator persistence. To this end, we numerically solve Eqs. 23 by varying the colonisation probability of the inferior prey (*c*_*y*_), which can enhance its performance relative to the superior prey, as well as the predator extinction probability (*e*_*z*_). Fig. 4 presents a phase diagram summarizing the different steady states obtained. For this purpose, a species was considered present when its stationary density exceeded a threshold of 0.0001. This representation shows in a single framework how coexistence patterns emerge from the combined effects of colonisation and extinction dynamics. In particular, it highlights the conditions under which the inferior prey can recover from competitive exclusion or when the predator’s persistence is compromised.

Let us first analyse Fig. 4 for *e*_*z*_ = 0, where the steady state depends on the colonisation probability of the inferior prey, *c*_*y*_. For *e*_*z*_ = 0 and *c*_*y*_ ≤ 0.4, the system reaches an absorbing state composed exclusively of predators. Indeed, once the last patch occupied by prey becomes empty, the dynamics freezes, since predator colonisation requires the presence of prey (see Eqs. 6–8). The predator density at these absorbing states depends on the initial conditions (figure not shown). This finding aligns with the infinite solutions identified for the steady states of Eqs. 24, which were discussed previously. When *c*_*y*_ is sufficiently large (*c*_*y*_ *>* 0.4), the system undergoes a transition to a coexistence phase between the inferior prey and the predator (*Y Z* steady-state). This result shows that the inferior prey can avoid extinction thanks to its high colonisation ability. This outcome is consistent with the fact that, for the set of parameters used in Fig. 4, species *Y* holds two key advantages over *X*: a higher colonisation probability and a lower extinction rate (see Table I). These traits compensate for its substantially higher predation probability. As a consequence, *X* does not persist because its limited colonising capacity, together with its higher extinction rate, dominates the benefit of being less susceptible to predation.

In the steady state corresponding to the *Y Z* coexistence for *e*_*z*_ = 0, the predator reaches its maximum density (*Z* = 1, figure not shown), since the persistence of *Y* ensures that all available patches can be colonized by *Z*.

Next, we analyse Fig. 4 for *e*_*z*_ *>* 0. For low values of *c*_*y*_, the system exhibits coexistence between the superior prey and the predator (*XZ* steady-state) at low *e*_*z*_. As *e*_*z*_ increases, the predator cannot persist, leaving a single-species state with only *X* (for *c*_*y*_ ≤ 0.06 ≈ 1.2*c*_*x*_). These results indicate that *Y* requires a minimum colonisation probability to survive alongside *X*, independent of predator presence. From this value of *c*_*y*_ upwards (*c*_*y*_ *>* 0.06 ≈ 1.2*c*_*x*_), the system enters a three-species coexistence phase (*XY Z* steady-state) at intermediate *e*_*z*_. For higher *e*_*z*_, when the predator is absent, the steady state shifts to coexistence between the two prey species (*XY*). Finally, for *c*_*y*_ *>* 0.14 ≈ 2.8*c*_*x*_, a *Y Z* coexistence phase emerges at low *e*_*z*_, indicating that the inferior prey can clearly outperform the superior prey under these conditions. The density maps used to construct Fig. 4 are presented in Appendix C.

**FIG. 4.**
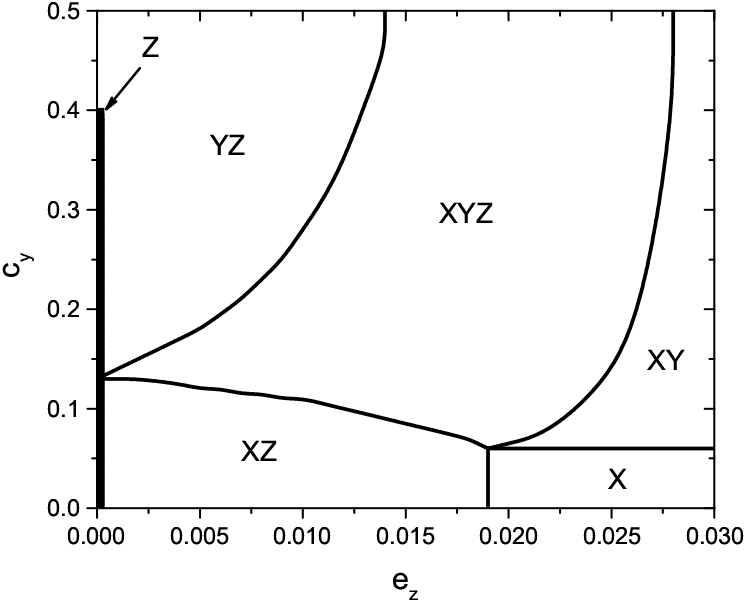
Mean-field phase diagram showing the steady-state of the model as a function of predator extinction probability *e*_*z*_ and inferior-prey colonisation probability *c*_*y*_ . Different steady states are indicated as *XY Z, Y Z, XZ, XY*, *X* and *Z*. The *Z* steady-state is found only for *e*_*z*_ = 0 and *c*_*y*_ ≤ 0.4. Parameter values are listed in Table I.

## V. STOCHASTIC SIMULATIONS

We performed stochastic simulations of our metapopulation model, bearing in mind that a patch occupied by a species has occupancy 1, while an uninhabited patch has occupancy 0. Furthermore, each patch can be occupied by one, two or three species at a time. The simulations were performed on a square lattice of size *L*, with a total of Ω = *L* × *L* patches. We found that for *L* ≥ *L*_min_ = 40, the steady-state properties no longer depend on lattice size; thus, we consider *L > L*_min_ in our study. In all Monte Carlo simulations, initial conditions were assigned randomly, with 50% of patches occupied by species *X* and *Y*, and 30% by species *Z*. The time to reach steady state varies with both lattice size and the parameter set. Closed boundary conditions were applied, and time was measured in Monte Carlo steps (MCS). At each MCS, the total number of patches is randomly chosen and the probability of the configuration update is evaluated in each case according to the definitions of Section II. The average fraction of patches occupied by each species is computed after the transient period.

First, let us analyse how the fraction of occupied patches for each species evolves over time for two different values of the predator extinction probability, as shown in Fig. 5. The system exhibits damped oscillations for the lower value of *e*_*z*_, which are more pronounced and longer-lasting than in the mean-field approach (shown in Fig. 3). An increase in prey density triggers a subsequent increase in predator density, which in turn reduces prey density, in a feedback loop. Also, there is a time delay between the maximum value of predator density and the minimum value of prey density, as observed in other predator-prey models [24, 26–28]. The steady-state for the lower value of *e*_*z*_ corresponds to coexistence between the inferior prey and the predator (*Y Z* steady-state), with most patches occupied by predators due to the low predator extinction probability. As *e*_*z*_ increases, the system reaches a three-species coexistence (*XY Z* steady-state), with the superior prey predominating after a transient period. In this case, typical stochastic fluctuations are observed.

**FIG. 5.**
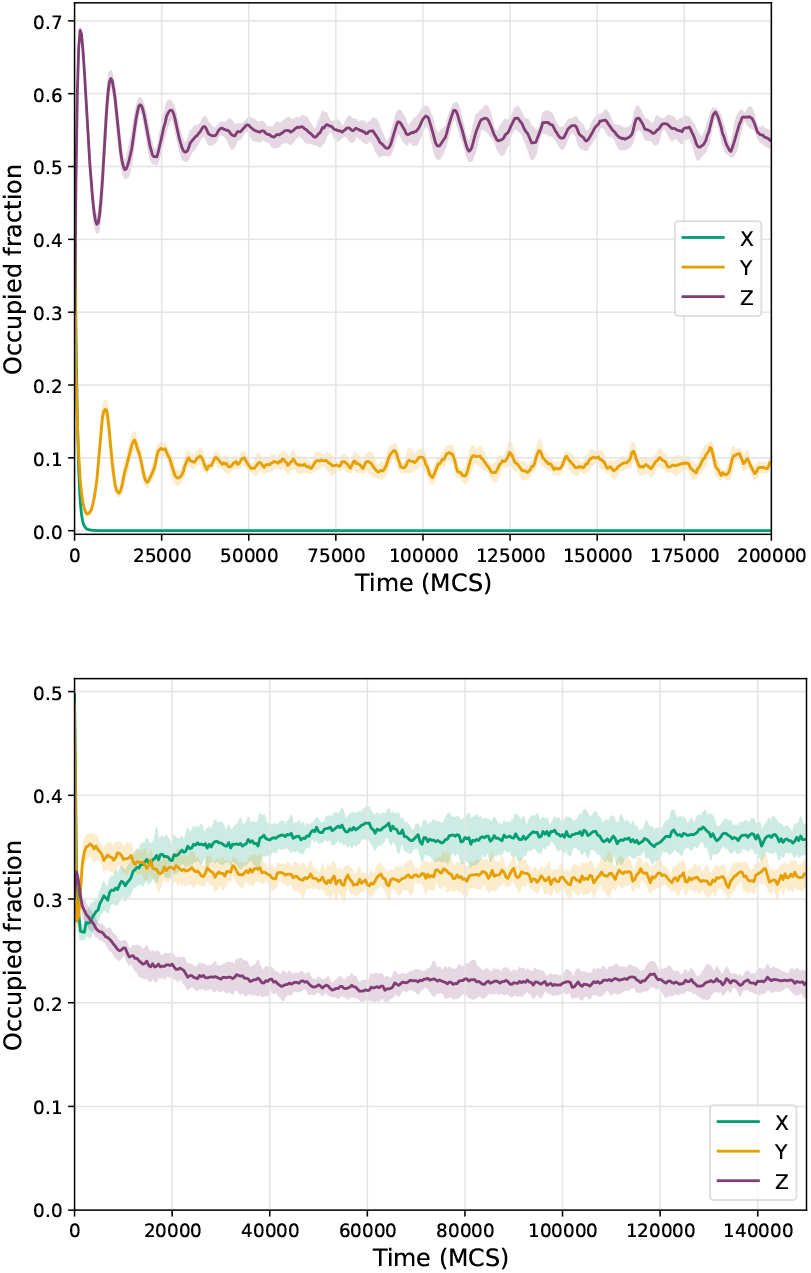
Time evolution of the fraction of patches occupied by species *X, Y*, and *Z* in stochastic simulations. Results correspond to parameters in Table I with *c*_*y*_ = 0.10 and *e*_*z*_ = 0.002 in the upper panel and *e*_*z*_ = 0.012 in the lower one. Solid lines represent the mean over 20 independent runs, and shaded regions indicate the corresponding standard deviation. Lattice size: *L* = 100.

Figure 6 shows the fraction of occupied patches at steady state as a function of the predator extinction probability, *e*_*z*_. When *e*_*z*_ = 0, the system reaches the absorbing state composed only of predators (*Z* steady state). For very small but non-zero values of *e*_*z*_, simulations become challenging, as the system can eventually get trapped in the trivial absorbing state, where *x* = *y* = *z* = 0. In fact, for the considered lattice sizes, the high fraction of patches occupied by predators can drive the prey species to extinction. This, in turn, leads to the extinction of the predators, since *e*_*z*_ is small but non-zero. The same lattice-size effect was also observed in other stochastic predator-prey model [24]. Besides, according to our analytical analytical results for the meanfield equations, we did not expect the trivial steady state to be stable for the parameters values considered here, as discussed in Section IV.

**FIG. 6.**
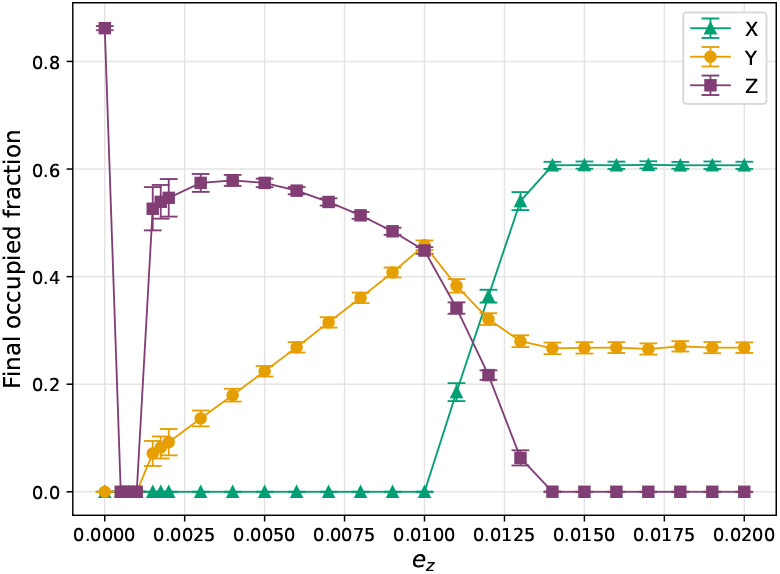
Mean values of the fraction of occupied patches as a function of the predator extinction probability, *e*_*z*_, for species *X* (superior prey), *Y* (inferior prey) and *Z* (predator), as obtained from Monte Carlo simulations at the stationary state. Mean values and standard deviations were calculated over the last 10^4^ points of the steady state, averaged across 20 independent runs of 2 × 10^5^ MCS each, for a system size *L* = 100. The parameters used are given in Table I, with *c*_*y*_ = 0.10.

As *e*_*z*_ increases, a sequence of state transitions is observed in the stochastic simulations. Initially, coexistence is established between the predator and the inferior prey (*Y Z* steady state). For intermediate values of *e*_*z*_, a threespecies coexistence emerges (*XY Z* steady state). Finally, at high values of *e*_*z*_, the predator goes extinct, leaving a two-prey coexistence (*XY* steady state).

When comparing Fig. 6 with the mean-field results in Fig. 2, it can be seen that the three-species coexistence and two-prey coexistence phases occur at intermediate and high *e*_*z*_, respectively, in both cases. The three-species coexistence region is more extended for meanfield (0.013 *< e*_*z*_ *<* 0.024) than for Monte Carlo simulations (0.01 *< e*_*z*_ *<* 0.014). However, the most striking finding is the qualitative difference observed at low *e*_*z*_: in the stochastic simulations, the predator coexists with the inferior prey (*Y Z* steady state), whereas in the meanfield approach, coexistence is between the predator and the superior prey (*XZ* steady state). Moreover, for low and intermediate values of *e*_*z*_, the relationship between the two prey species differs significantly between the two methodologies. In mean-field, the superior prey always maintains a higher density than the inferior prey, while in Monte Carlo simulations, the inferior prey presents an advantage in relation to the superior prey only when predator coverage is high. To understand this qualitative difference between mean-field and Monte Carlo results, let us analyse the spatial distribution of the species.

Figure 7 shows the spatial distribution of species in the steady state for a three-species coexistence scenario (*XY Z* steady-state). Panel (a) displays each species separately, panel (b) shows all species together in the same lattice, while panel (c) highlights only the coexistence patches. It is evident from Fig. 7-(a) that the model presents a spatial pattern. In fact, each species tends to form clusters, due to the model rules in which colonisation occurs from already occupied patches. In Fig. 7-(b), predators are observed in the vicinity of prey patches, since their spreading depends on them according to the rules of colonisation.

**FIG. 7.**
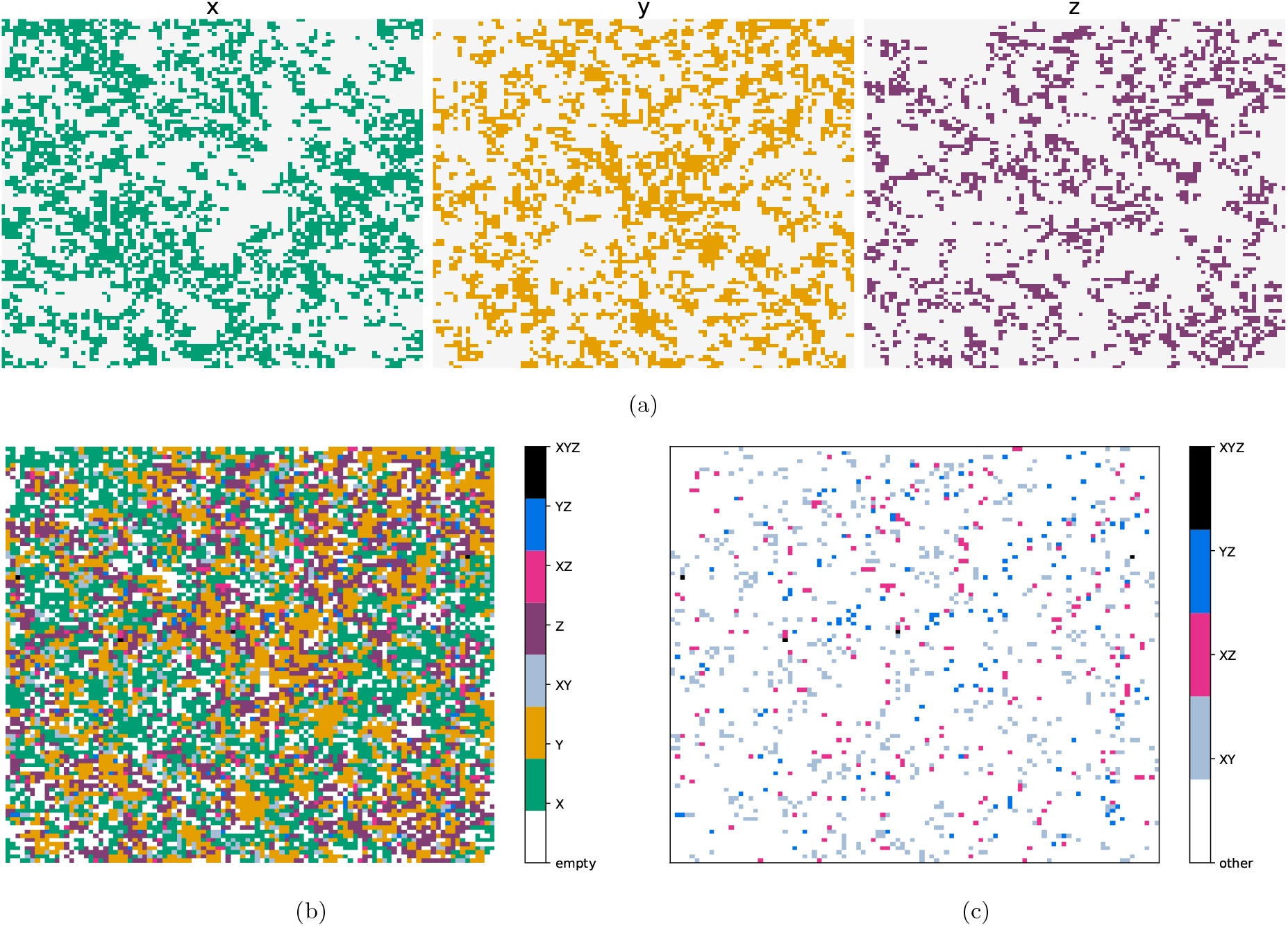
Spatial distribution of species in the steady-state (*XY Z* steady-state). (a) Distribution of each species individually: *X* (left), *Y* (centre), and *Z* (right). Unoccupied patches are shown in light grey. (b) All species together in a single lattice, with different colours indicating coexistence patches. Empty patches are shown in white. (c) Coexistence patches only; empty patches and patches occupied by a single species are shown in white, while coloured patches indicate coexistence. The parameters used are given in Table I, with *c*_*y*_ = 0.10 and *e*_*z*_ = 0.012. Simulation details: *MCS* = 2 × 10^5^, *L* = 100.

Patches where coexistence occurs are much less frequent than patches occupied by a single species, as is evident from Fig. 7-(c). In particular, coexistence between the two prey (*XY* patches ≈ 6%) is more common than coexistence of prey with the predator (*XZ* ≈ 2%, *Y Z* ≈ 1.5%) or with both prey and the predator (*XY Z* ≈ 0.04%). One possible reason for these results is that, given the parameters considered, the probabilities of *X* and *Y* being preyed upon is four and sixteen times greater, respectively, than the probability of *Y* being eliminated due to hierarchical competition with *X*. Overall, the results of Fig. 7 shows a spatial distribution in which single-species patches dominate, *XY* coexistence is more common than coexistence involving the predator, and the spatial patterns reflect the combined effects of colonisation, predation, and hierarchical competition.

From these results, we can conclude that spatial aggregation plays an important role in the model. When the spatial coverage of predators is high, predation becomes a dominant process, and prey colonisation is crucial for their persistence. In such situations, the spatial aggregation of prey is particularly relevant: when forming clusters, patches in the centre of a cluster are protected from predation and are maintained over time. Although the predation probability on *Y* is four times higher than on *X*, the inferior prey has twice the colonisation probability of the superior prey. The results of Fig. 6 indicate that, in the stochastic spatial model, the colonisation advantage can outweigh the predation disadvantage. Consequently, for low values of *e*_*z*_, coexistence occurs between the predator and the inferior prey.

In contrast, the spatial aggregation of prey is not captured by our mean-field equations, which neglect spatial correlations. In this framework, predation dominates at low *e*_*z*_, giving the superior prey an advantage over the inferior prey. As a result, the mean-field model predicts coexistence between the predator and the superior prey, as illustrated in Fig. 2.

## VI. DISCUSSION

There are many ways to formulate quantitative models in ecology. One popular methodology is the use of phenomenological mean-field equations, which are proposed on an *ad hoc* basis. These equations are simple to construct and implement. Another option is to propose individual-based models, which offer a more accurate description of the system as they take into account prior knowledge about individuals or species and their interactions. Computer simulations are typically employed for individual-based models, and their implementation can be slightly more challenging than numerical resolution of mean-field equations. A third option is to derive first-principles mean-field equations from the master equation of the individual-based model. In this case, a well-defined approximation scheme is employed and comparison with Monte Carlo simulation results is straightforward. One reason for this is that, when working with first-principles mean-field equations, the relationship between the parameters used in both methodologies can be determined analytically. Analysing stationary states and their stability using theoretical and numerical tools can provide a better understanding of the simulation results.

In this paper, we combine first-principles mean-field equations and stochastic simulations. We examine a spatially explicit three-species metapopulation model consisting of one predator and two prey species. We consider the biological processes of extinction, interspecific competition between prey, predation, and colonisation. We take into account specific rules for colonisation that represent the predator’s search for prey and the hierarchical competition among prey. We derive the master equation for the model and obtain a mean-field approximation for the time evolution of the mean occupancy of each species. We also perform stochastic simulations to explore the system’s behaviour beyond the mean-field level.

We discuss that our first-principle mean-field equations are quite simple and would even be naively write down on an *ad hoc* basis. However, in the case of a model with complex colonisation rules, it can be difficult to write the correct mean-field terms without a first-principles strategy. In fact, other authors have proposed *ad hoc* equations for the same [4] or similar system [19], that differ from our first-principle equations. These differences ultimately translate into quantitative and sometimes qualitative changes in the predicted stability and coexistence domains. Besides, when deriving mean-field equations from the master equation, a well-defined approximation scheme is used, and different levels of approximations can be made [1, 21, 24, 25]. Moreover, the relationship between the model parameters and the mean-field parameters is univocally determined only with a first-principles formulation.

A steady-state analysis of the first-principle mean-field equations shows that the system can exhibit up to eight steady-states, corresponding to the different coexistence combinations among the three species. By linearising the equations about the steady-states we find that the system is always monostable, and that different steadystate solutions alternate as parameters vary. The stable fixed points are classified as either stable nodes or spiral nodes. As expected, numerical solutions of the mean-field equations display damped oscillations for the parameter values where spiral nodes are found. These oscillations appear at low values of the predator extinction probability. For two-species predator-prey models, oscillatory behaviour is also observed at low values of the predator extinction probability, although in those cases sustained oscillations have been reported [24, 26, 29]. We guess that the more complex dynamics involving three interacting species in our model prevents the instability of the fixed points, and therefore, the presence of oscillations.

In our model, Monte Carlo results present a more pronounced damped oscillations than the mean-field results, probably because stochasticity is an additional source of non-linearity that can contribute to oscillatory behaviour. Indeed, some stochastic models exhibit sustained oscillations that do not appear in the deterministic population-level description, a phenomenon known as noise-induced oscillation [30, 31].

We observe qualitative agreement between our meanfield results and Monte Carlo simulations for both the three-species coexistence and the coexistence between the two prey. However, the mean-field description fails to reproduce the coexistence between the predator and the inferior prey observed at low predator extinction probabilities. This discrepancy arises from the neglect of spatial correlations in the mean-field approach: in the stochastic model, spatial clustering of the inferior prey provides effective protection against predation. The comparison between mean-field and Monte Carlo results provides an opportunity to improve our understanding of the relationship between colonisation, predation and spatial aggregation in our model. These results underscore the importance of combining complementary modelling approaches, even for relatively simple ecological systems [32].

Beyond its methodological relevance, our results emphasise that even minimal spatial structure can generate ecological outcomes that differ qualitatively from those predicted by well-mixed approximations. In particular, the coexistence between the predator and the inferior prey, absent in the mean-field description yet recovered in the stochastic formulation, underscores the crucial role of spatial correlations and local interactions in shaping community dynamics. These findings suggest that, in ecological systems where colonisation and predation act on comparable spatial scales, aggregation effects can act as a stabilising mechanism that promotes coexistence.

Overall, the first-principles framework developed here provides a systematic bridge between individual-based dynamics and population-level descriptions. It enables analytical insight into the correspondence between stochastic and deterministic formulations, while exposing the limits of mean-field approximations. Extending this approach to include explicit spatial coupling, higherorder correlation closures, or additional trophic levels would offer promising directions for future work, contributing to a deeper theoretical understanding of spatial coexistence in ecological metacommunities.

## VII APPENDIX A

In this appendix, we present an alternative approach to deriving the mean-field equations, starting from Eqs. 22 in Section III. As in Section IV, we assume that patches on the grid are spatially uncorrelated, that is, they are independent from one another. Therefore, we consider ⟨*X*_*i*_*Y*_*i*_⟩= ⟨*X*_*i*_⟩ ⟨*Y*_*i*_⟩, and similar for other mean values in Eq. 22. Under this assumption, for species *X* at patch *i* we obtain:

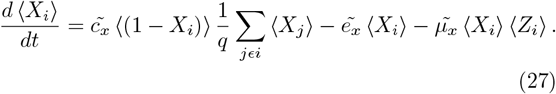

If we add and subtract the quantity 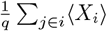 in Eq. 27, we obtain the discrete Laplacian, defined as:

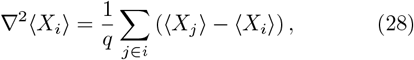

with unit lattice spacing. After that, we can assume that all sites are equivalent, ⟨*X*_*i*_⟩ = ⟨*X*_*j*_⟩ = ⟨*X*⟩ = *x*, and that they are distributed in an infinite domain, i.e., Ω → ∞. Under these assumptions, and with a similar procedure for species *Y* and *Z*, we obtain the following first-principle mean-field equations:

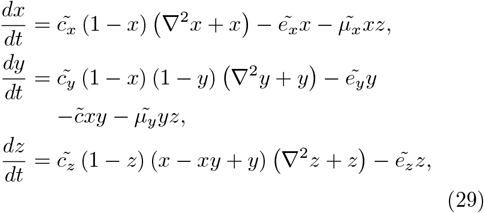

where the Laplacian in each equation accounts for the diffusive contribution of colonisation.

To study the system’s dynamics, we numerically integrated the spatially explicit mean-field equations (Eqs. 29) in two-dimensions. The spatial component was integrated using the finite difference approximation for the second derivative (Eq. 28), and for the temporal component, we employed the second-order Runge–Kutta method. We considered a square lattice of size *L* = 50, corresponding to Ω = 250 patches. The initial distributions of *X, Y*, and *Z* were Gaussian and radially symmetric, each with a standard deviation of *σ* = 0.8. The peak of each Gaussian was located at the centre of the lattice, reaching a maximum value of 0.50 for all species. Von Neumann boundary conditions were applied, meaning that the derivative at the edges of the lattice vanishes, ensuring a closed system. The simulation was run for 10^5^ time steps.

In Fig. 8 we analyse how the initial spatial perturbation in density evolves. For this purpose, we selected the central row of the lattice, *i* = 25, and recorded *X, Y* and *Z* densities along this line, for several times. From the curves, we observe that the species gradually spread across the lattice until they reach a homogeneous distribution. This indicates that no spatial patterns are observed at the steady state, despite the presence of the initial inhomogeneity.

**FIG. 8.**
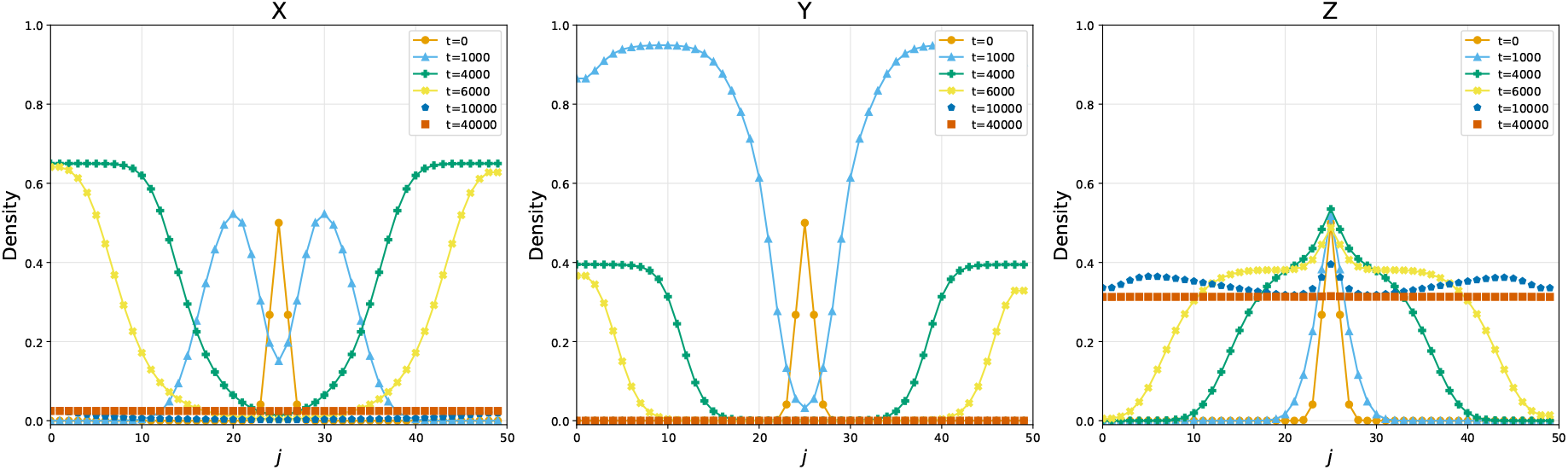
Density profiles of species *X* (left), *Y* (centre), and *Z* (right), for the spatial mean-field model at several simulation times. The profiles corresponds to the middle row (*i* = 25) of a lattice of size *L* = 50. The system reaches the *XZ* coexistence steady state. The parameters are listed in Table I, with *c*_*y*_ = 0.10 and *e*_*z*_ = 0.0005 (the same as those used in the upper panel of Fig. 3).

A comparison between the spatial and non-spatial mean-field models is illustrated in Fig. 9, which displays the time evolution of the densities of *X, Y*, and *Z*. Both correspond to *e*_*z*_ = 0.0005 and the parameters listed in Table I. Although the models differ during the transient regime, they converge to the same steady state, the *XZ* coexistence. Damped oscillations are observed in both the spatial and non-spatial models for this value of *e*_*z*_.

**FIG. 9.**
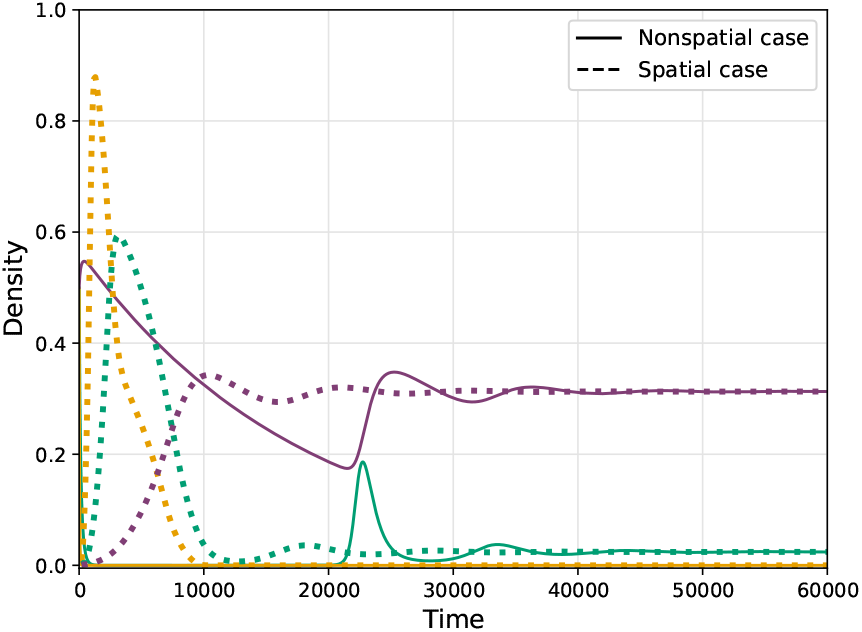
Mean-field results for the time evolution of the densities of species *X* (green), *Y* (yellow), and *Z* (purple) for the spatial (dotted lines) and non-spatial (solid lines) cases. The parameters are listed in Table I, with *c*_*y*_ = 0.10 and *e*_*z*_ = 0.0005 (the same as those used in Fig. 8 and in the upper panel of Fig. 3).

Since both models reach the same stationary state, the non-spatial formulation is sufficient for the mean-field description, being simpler and computationally more efficient. The same behaviour was observed for other values of *e*_*z*_ and different initial conditions.

## VII. APPENDIX B

The elements of the Jacobian Matrix presented in Section II are given by

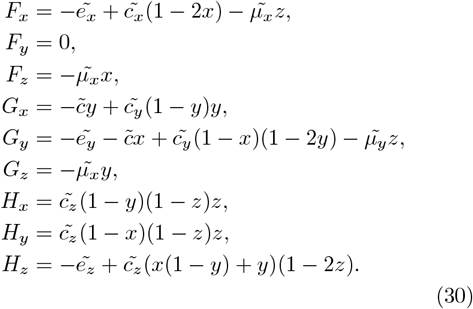

The secular equation *c*_0_ + *c*_1_*λ* + *c*_2_*λ*^2^ − *λ*^3^ = 0 (Eq. 26) presented in Section II, has the following exact coefficients

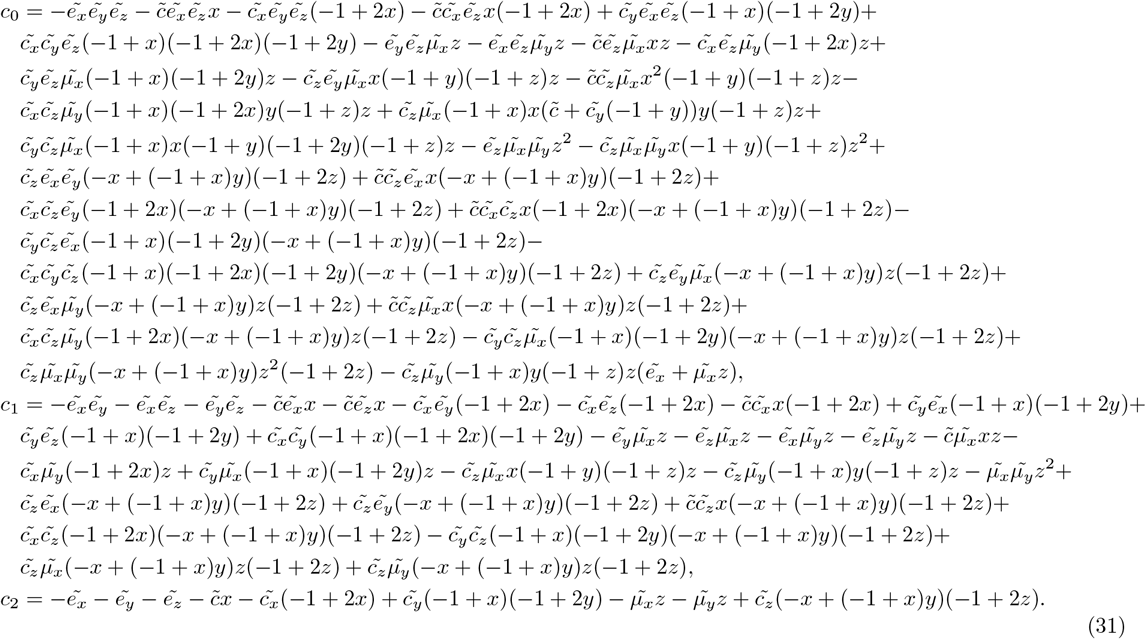

## IX. APPENDIX C

In this appendix, we present the density maps used to construct Fig. 4 in Section IV. Fig. 10 shows the steady-state densities reached by species *X, Y*, and *Z*, for different values of the extinction probability of the predator (*e*_*z*_) and the colonisation probability of the inferior competitor (*c*_*y*_). The intervals for *e*_*z*_ and *c*_*y*_ were chosen because, within these ranges, we observed significant changes in the system’s dynamics. From Fig. 10, we can observe how different parameter combinations promote either extinction or persistence of the species, as also shown in Fig. 4.

**FIG. 10.**
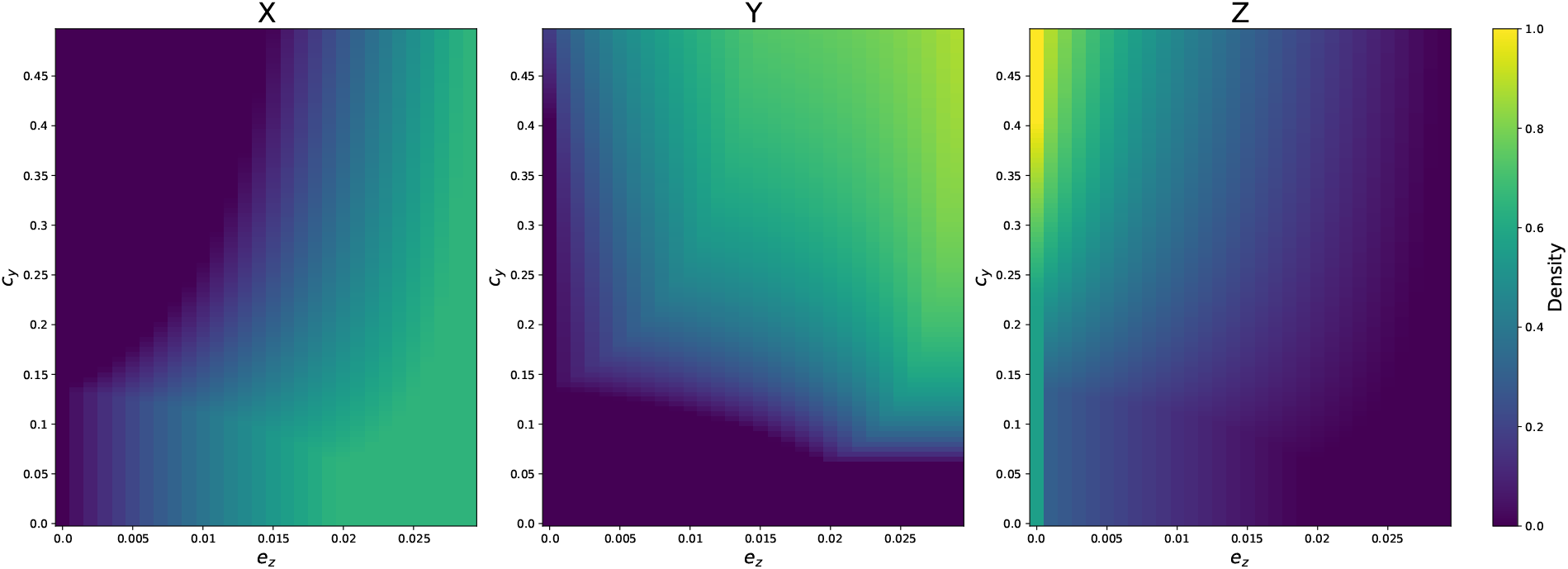
Steady-state density maps for species *X* (left), *Y* (centre) and *Z* (right) obtained from mean-field. The *e*_*z*_ and *c*_*y*_ steps were chosen as 0.001 and 0.005, respectively. Results correspond to 2 × 10^6^ MCS.

